# Systematic position of edible tropical oysters and first record of genus *Talonostrea* (Ostreidae) in the Myeik coastal area, Myanmar

**DOI:** 10.1101/2025.01.29.635588

**Authors:** Masaya Toyokawa, Tatsuya Yurimoto, Bong Jung Kang, Zakea Sultana, Cherry Aung

## Abstract

Belcher’s (*Magallana belcheri*) and hooded (*Saccostrea cuccullata*) oysters are edible species in Myeik, Myanmar. Belcher’s oyster is particularly desirable for aquaculture because of its high price and potential value as export products. Previous studies have noted a mismatch between oyster spat settlement patterns and the maturity of the species’ gonads, suggesting the possible settling of unintended oyster species in the Myeik coastal area. To address this issue, DNA barcoding was used to identify oyster species collected from the Myeik coast. Genetic analysis of mitochondrial DNA sequences from various sources of oysters sampled from mangrove roots, experimental cultivation rafts, and settled natural spat revealed an abundance of unidentified species of the genus *Talonostrea* in addition to Belcher’s and hooded oysters. Therefore, this study suggested that collecting aquaculture target oysters from natural spat is inefficient because only a low percentage is found in coastal areas.

## Introduction

Belcher’s oyster *Magallana belcheri* (G. B. Sowerby II, 1871) and hooded oyster *Saccostrea cuccullata* (Born, 1778) are known as Ka-mar in Myanmar and are edible species (Yurimoto et al. 2019; Oo 2020, 2021; Khin-May-Chit-Maung et al. 2023). Because of the high demand for the larger edible oyster, *M. belcheri*, we began a study on oyster farming in the Myeik coastal area of southern Myanmar in 2017. Marine aquaculture of prawns, crabs, and fish is developing, but the aquaculture of bivalves is not mentioned by the Myanmar government in its review of aquaculture and resource enhancement (Aung 2021). The exception is the highly developed pearl culture industry, which exported an average of 13.7 million USD annually from 2015 to 2020, according to the United Nations Commodity Trade Statics Database (http://comtradeplus.un.org/db/ [Accessed on 18 March 2024]). This amount is comparable to 2% of the total exports of fish and fishery products, which was 712 million USD in the 2017–2018 fiscal year (Aung 2021).

One of the keys to developing oyster farming is to provide seeds for grow-out cultures. In most tropical countries, oyster spat is sourced from wild collections (Nowland et al. 2020). To effectively collect oyster seeds from the wild, it is essential to determine the peak settling season of the target species. Although gonads in the spawning stage were observed at the highest rate in October–November and April–May for *M. belcheri* in the Myeik coastal area (Khin-May-Chit-Maung et al. 2023), the highest number of settled spat was observed in different months, depending on the station (Aung-Aung-Aye et al. unpublished data). Previous studies on oyster spat collection in Myanmar have identified oyster spat species from the perspective of shell morphology. For example, Thi-Thi-Lay (1983) classified spat with a white or black stripe from the hinge to shell edges as *S. cuccullata*. However, during our study of oyster seed collection from the wild, we noticed that it was difficult to identify the species of settled oyster seeds based on their appearance. Therefore, we decided to analyse the DNA and identify the species genetically.

Genetic identification of marine species is not yet well established in Myanmar. Specimens must be exported to other countries to analyse DNA sequences through a material transfer agreement (MTA). Through a genetic study of oysters carried out under MTA between Myeik University and the Japan International Research Center for Agricultural Sciences (JIRCAS), we recognised that a considerable proportion of the collected oysters belonged to the genus *Talonostrea*. *Talonostrea* oysters have never been reported on the coast of Myanmar. Therefore, we report the preliminary results of a genetic study with the first record of the genus *Talonostrea* in Myanmar.

## Materials and methods

### Sample collection and DNA sequencing

Cultured oysters were sampled from an experimental oyster culture cage, and wild oysters were collected from a mangrove root near the oyster cage in Pa-yi-kyun village on 12 November 2019 (Figure 1). Oyster spat samples were collected from spat collectors set in a fish cage in a creek near Pe-taing village from 5 February to 5 March 2020. Samples were transported to the laboratory at Myeik University and preserved in ethanol.

**FIGURE 1.**
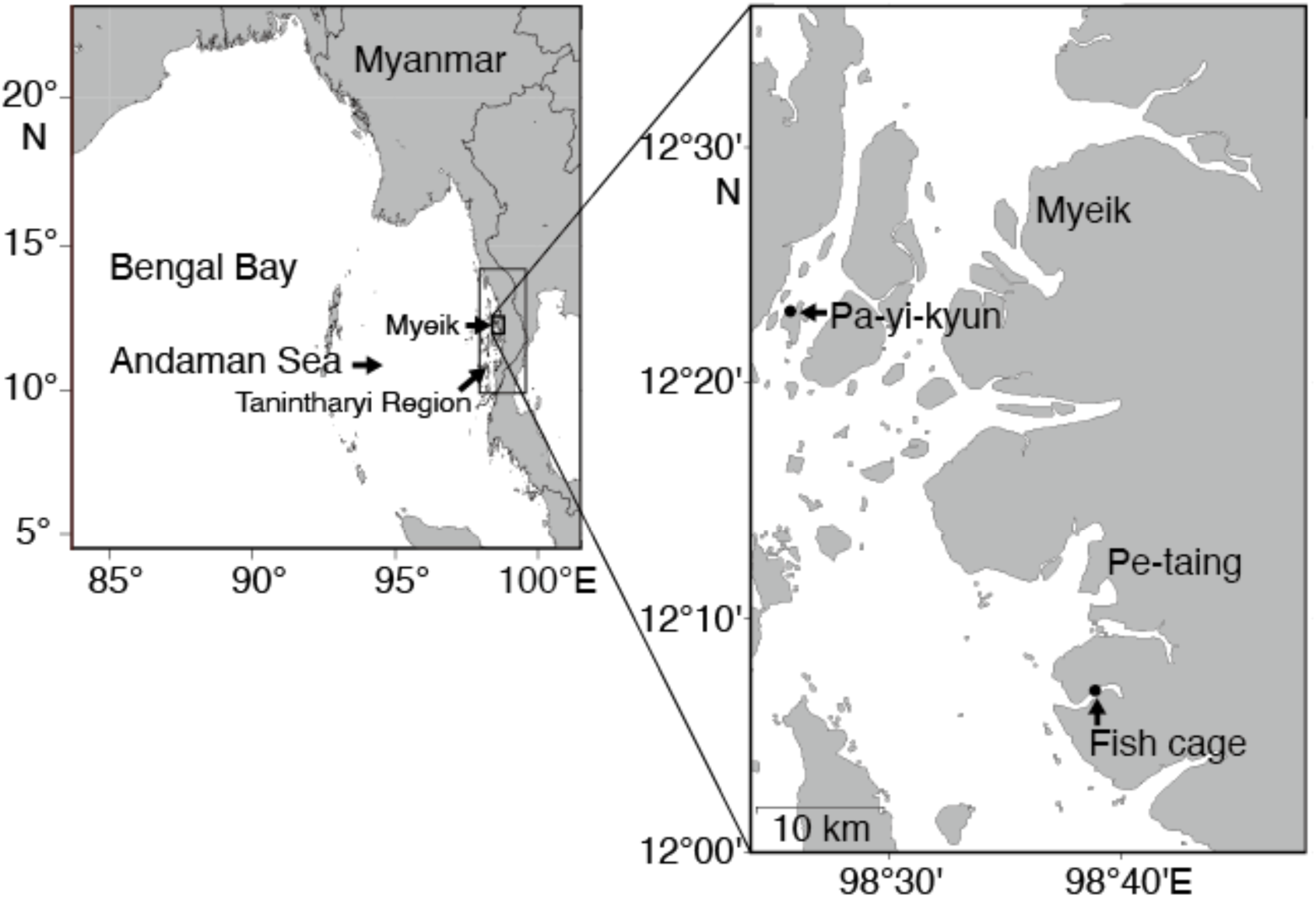
Map showing the sample collection site of the present study in Pa-yi-kyun village and a fish cage near Pe-taing village.

Shell length, height, and width were measured using a digital calliper (CD-S20C; Mitsutoyo Co. Ltd., Kanagawa, Japan), and weights with and without shells were determined using a mini-digital scale (DS-22; Juanjuan Electronic Technology Co., Ltd., Guangzhou, China). Tissues were dissected from the ethanol-preserved samples. DNA was extracted and purified using a DNeasy 96 Blood and Tissue Kit (QIAGEN, Hilden, Germany) following the manufacturer’s instructions. Partial sequences of two mitochondrial DNA regions, cytochrome c oxidase subunit I (COI) and 16S *rRNA* (16S) gene were amplified using PCR with universal COI primers, LCO1490 (5ʹ-GGT CAA CAA ATC ATA AAG ATA TTG G-3ʹ) and HCO2198 (5ʹ-TAA ACT TCA GGG TGA CCA AAA AAT CA-3ʹ), and with 16S primers, 16Sar (5ʹ-CGC CTG TTT ATC AAA AAC AT-3ʹ) and 16Sbr (5ʹ-CCG GTC TGA ACT CAG ATC ACG T-3ʹ). The PCR cycling parameters were 94°C for 4 min followed by 32 cycles of 94°C for 20 s, 56°C for 40 s and 72°C for 1 min, and a final extension at 72°C for 1 min. PCR products were sequenced by Fasmac DNA sequencing service (FASMAC, Atsugi, Japan) using the same primer sets as those used for PCR.

Each sequence result was checked for the ABI quality value (QV) using the Applied Biosystems DNA Sequencing Analysis software manual (Applied Biosystems 2003), and the base was replaced with X where QV was less than 15. The start and end parts of a sequence, with low (<15) QV, were trimmed. Samples with a generally low QV were deleted. Sequences that cleared the screening process were further screened using a BLAST search at the National Center for Biotechnology Information (NCBI). Sequences that probably resulted from contamination or misidentification were excluded from the analysis. After the removal of short-length sequences (<480 for COI or <400 for 16S), the remaining sequences were used in the subsequent phylogenetic analyses.

### Phylogenetic analyses

The screened sequences were classified into two groups (Crassostreinae and *Saccostrea*) based on the BLAST search results. The two groups were analysed separately using reference sequences (Table 1). The reference sequences of Li et al. (2017a) and Sekino and Yamashita (2016) were followed. Sequences (COI or 16S) of *Magallana* and *Talonostrea* that were reported since 2017 and sequences of valid *Crassostrea* species listed in the World Register of Marine Species (WoRMS) were included in the reference sequences as long as the sequences were retrieved from the nucleotide database of the NCBI. The sequences of specimens A–F, Y, and Z from Li et al. (2017a) which were collected from the Tanintharyi Region of Myanmar, were included in the analyses for comparison (see Table 1 of Li et al. 2017a). Alignment was performed using MAFFT v7.490 using the L-INS-i strategy (Katoh and Standley 2013). For COI, the DNA sequences were translated into amino acid sequences using pgtransec (Tanabe 2008) and the amino acid sequences were aligned using MAFFT. The aligned sequences were translated back into DNA sequences. Aligned sequences were trimmed manually to obtain the same length and automatically trimmed using trimAl to remove ambiguously aligned segments (Capella-Gutiérrez et al. 2009). Duplicate sequences were grouped together and treated as haplotypes.

**Table 1.**
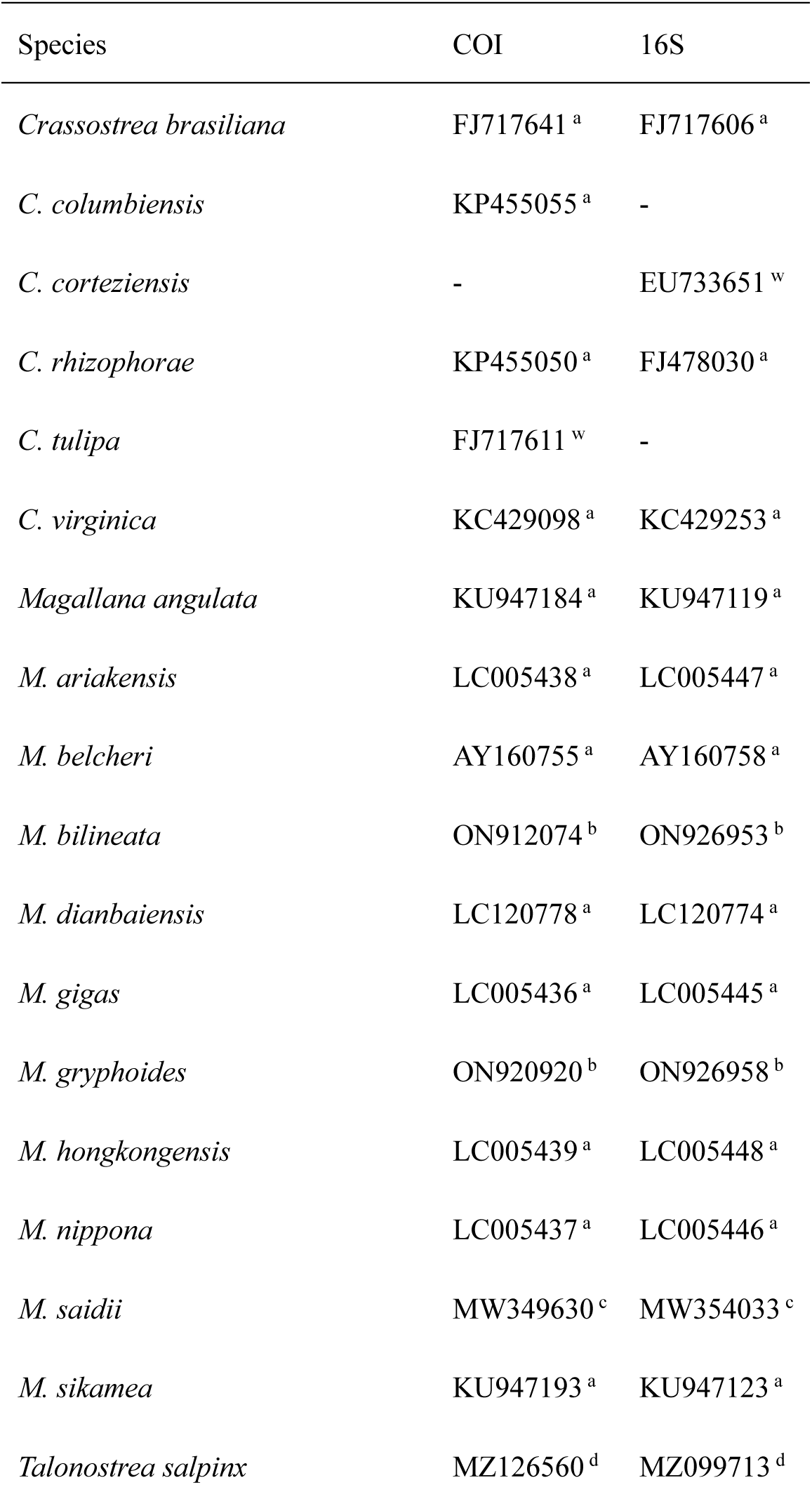
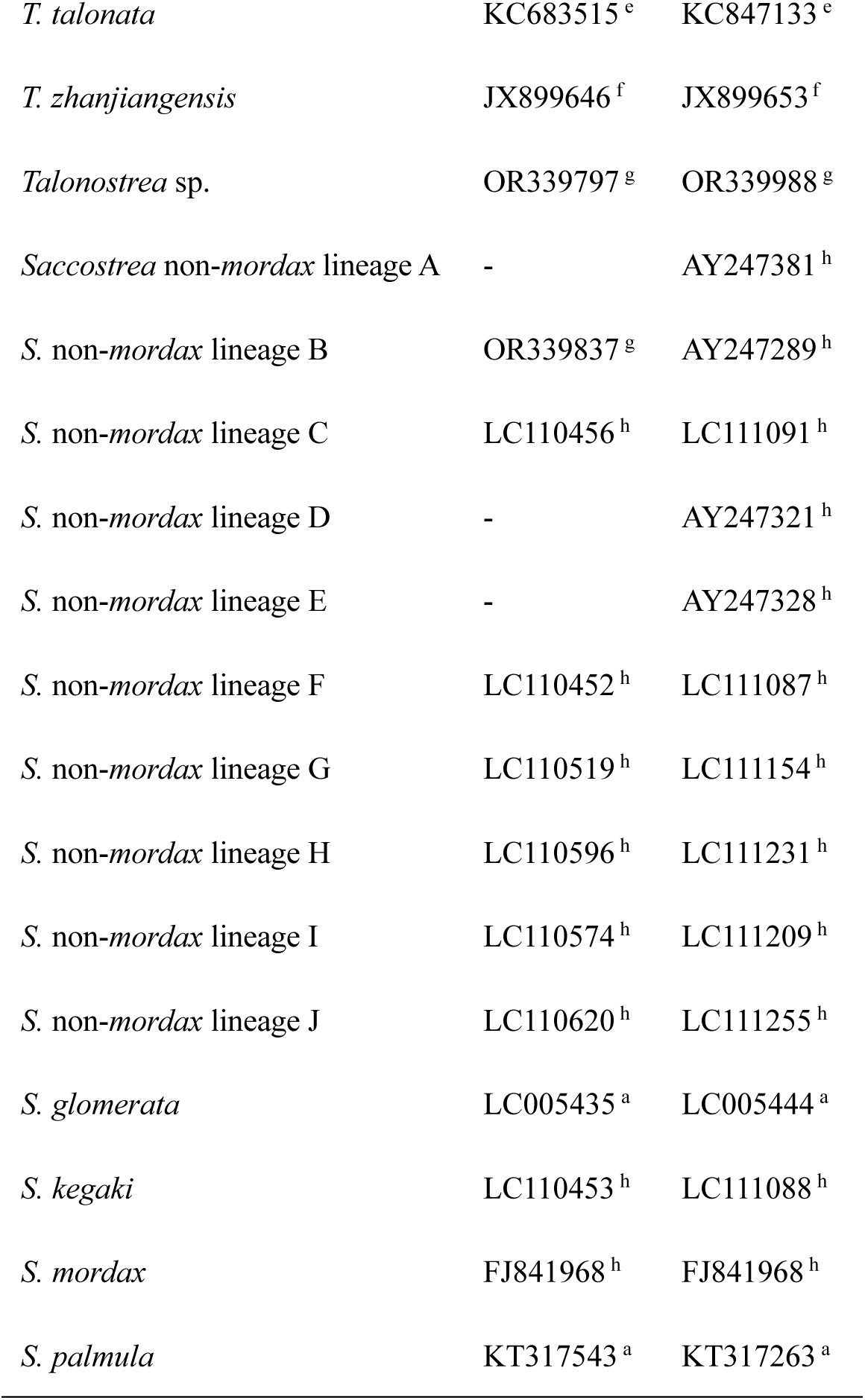
Reference DNA sequences of oysters which were taken from the reference lists of previous taxonomic studies. a: Li et al. (2017a); b: Santhi et al. (2022); c: Sigwart et al (2021); d: Al-Kandari et al. (2021); e: Salvi and Mariottini (2017); f: Wu et al. (2013); g: McDougall et al. (2024); h: Sekino and Yamashita (2016); w: valid species on the WoRMS and sequences were retrieved from the NCBI nucleotide database. – denotes that sequence was not available from the NCBI nucleotide database. NA denotes that location is not available.

| Species | COI | 16S |
| --- | --- | --- |
| *Crassostrea brasiliana* | FJ717641<sup>a</sup> | FJ717606<sup>a</sup> |
| *C. columbiensis* | KP455055<sup>a</sup> | - |
| *C. corteziensis* | - | EU733651<sup>w</sup> |
| *C. rhizophorae* | KP455050<sup>a</sup> | FJ478030<sup>a</sup> |
| *C. tulipa* | FJ717611<sup>w</sup> | - |
| *C. virginica* | KC429098<sup>a</sup> | KC429253<sup>a</sup> |
| *Magallana angulata* | KU947184<sup>a</sup> | KU947119<sup>a</sup> |
| *M. ariakensis* | LC005438<sup>a</sup> | LC005447<sup>a</sup> |
| *M. belcheri* | AY160755<sup>a</sup> | AY160758<sup>a</sup> |
| *M. bilineata* | ON912074<sup>b</sup> | ON926953<sup>b</sup> |
| *M. dianbaiensis* | LC120778<sup>a</sup> | LC120774<sup>a</sup> |
| *M. gigas* | LC005436<sup>a</sup> | LC005445<sup>a</sup> |
| *M. gryphoides* | ON920920<sup>b</sup> | ON926958<sup>b</sup> |
| *M. hongkongensis* | LC005439<sup>a</sup> | LC005448<sup>a</sup> |
| *M. nippona* | LC005437<sup>a</sup> | LC005446<sup>a</sup> |
| *M. saidii* | MW349630<sup>c</sup> | MW354033<sup>c</sup> |
| *M. sikamea* | KU947193<sup>a</sup> | KU947123<sup>a</sup> |
| *Talonostrea salpinx* | MZ126560<sup>d</sup> | MZ099713<sup>d</sup> |
| *T. talonata* | KC683515<sup>e</sup> | KC847133<sup>e</sup> |
| *T. zhanjiangensis* | JX899646<sup>f</sup> | JX899653<sup>f</sup> |
| *Talonostrea* sp. | OR339797<sup>g</sup> | OR339988<sup>g</sup> |
| *Saccostrea non-mordax* lineage A | - | AY247381<sup>h</sup> |
| *S. non-mordax* lineage B | OR339837<sup>g</sup> | AY247289<sup>h</sup> |
| *S. non-mordax* lineage C | LC110456<sup>h</sup> | LC111091<sup>h</sup> |
| *S. non-mordax* lineage D | - | AY247321<sup>h</sup> |
| *S. non-mordax* lineage E | - | AY247328<sup>h</sup> |
| *S. non-mordax* lineage F | LC110452<sup>h</sup> | LC111087<sup>h</sup> |
| *S. non-mordax* lineage G | LC110519<sup>h</sup> | LC111154<sup>h</sup> |
| *S. non-mordax* lineage H | LC110596<sup>h</sup> | LC111231<sup>h</sup> |
| *S. non-mordax* lineage I | LC110574<sup>h</sup> | LC111209<sup>h</sup> |
| *S. non-mordax* lineage J | LC110620<sup>h</sup> | LC111255<sup>h</sup> |
| *S. glomerata* | LC005435<sup>a</sup> | LC005444<sup>a</sup> |
| *S. kegaki* | LC110453<sup>h</sup> | LC111088<sup>h</sup> |
| *S. mordax* | FJ841968<sup>h</sup> | FJ841968<sup>h</sup> |
| *S. palmula* | KT317543<sup>a</sup> | KT317263<sup>a</sup> |

The best models for phylogenetic analyses were selected using the Akaike information criterion corrected for small samples (AICC) using ModelTest-NG (Darriba et al. 2020; Flouri et al. 2014). The maximum likelihood (ML) tree with the selected model was inferred using RAxML-NG v. 1.1 (Kozlov et al. 2019). Branch support levels were calculated from the bootstrap values of 2000 replicates. The mean number of substitutions per site was expressed as branch length.

### Shell morphology

Four morphological characteristics of the shells of specimens identified using phylogenetic analyses were observed and recorded: the presence or absence of chomata, dark staining on the interior valve, radial ribs of the left valve, and a black or white streak on the outer surface of the right valve.

Images of the oyster shells (the insides and outsides of both shells) were captured using a Canon Camera Connect version 3.1 on an Apple iPad through a Canon 5D Mark IV with a Canon EF 100 mm f/2.8L Macro IS USM lens. Depending on the size of the shell, approximately 10–20 images were taken from the highest point of the shell to the lowest plane by moving the focus points step-by-step with the focus function of the Canon Camera Connect software. Then three-D focus stacked images were rendered from each sample using Helicon Focus version 8.2.10 Pro (Helicon Soft Ltd., Kharkiv, Ukraine). Each stacked image was trimmed, and the scale was replaced with a vector line representing 5 mm using Adobe Illustrator version 28 (Adobe Inc., San Jose, USA).

## Results

### Phylogenetic analyses

From a total of 137 specimens (42 from a mangrove root, 80 from cultured, and 15 from wild spat), 61 COI and 126 16S sequences were retrieved after quality control and BLAST searches (Table 2). These sequences were classified into either the subfamily Crassostreinae (COI: 36 sequences; 16S: 80 sequences), or the genus *Saccostrea* (COI: 25 sequences; 16S: 46 sequences) using a BLAST search on NCBI (Table 2). Oysters from other groups were not found in the samples. Removal of the short sequences resulted in 39 COI sequences and 115 16S sequences, which were grouped into 22 haplotypes for COI and 20 haplotypes for 16S (Table 3). Among the seven haplotypes of the 16S sequences of Crassostreinae oysters, five were in the same clade as *Talonostrea talonata* (Figure 2). One haplotype derived from ten specimens had a sequence identical to that of the reference *T. talonata* sequence (KC847133). The other two haplotypes formed a monophyletic group with the four *Talonostrea* species; however, they differed from all other *Talonostrea* species (Figure 2). Among the seven haplotypes of the COI sequences of the Crassostreinae oysters, a single haplotype represented by a single specimen formed a clade with *Magallana belcheri* (Figure 3). However, the 16S sequence was not retrieved from this specimen because of its quality. The other six haplotypes formed a monophyletic group with the four *Talonostrea* species and were different from all four *Talonostrea* species (Figure 3). Specimens belonging to the clade separated from other *Talonostrea* species were all derived from the root of a mangrove tree. The specimen sequences formed a distinct clade with a small genetic distance from each other. The expected substitution rate per site within the haplotypes in this clade was 0.00311 for the 16S ML tree and 0.00382–0.01745 for the COI ML tree. These values were considerably smaller than the distance between this group and one of the other four *Talonostrea* species (0.1317–0.1668 for 16S and 0.4434–0.6301 for COI). This distance was comparable to the distance between the other four *Talonostrea* species (0.0128–0.1260 for 16S and 0.1181–0.7455 for COI). Hereafter, this group is referred to as *Talonostrea* sp. 1.

**FIGURE 2.**
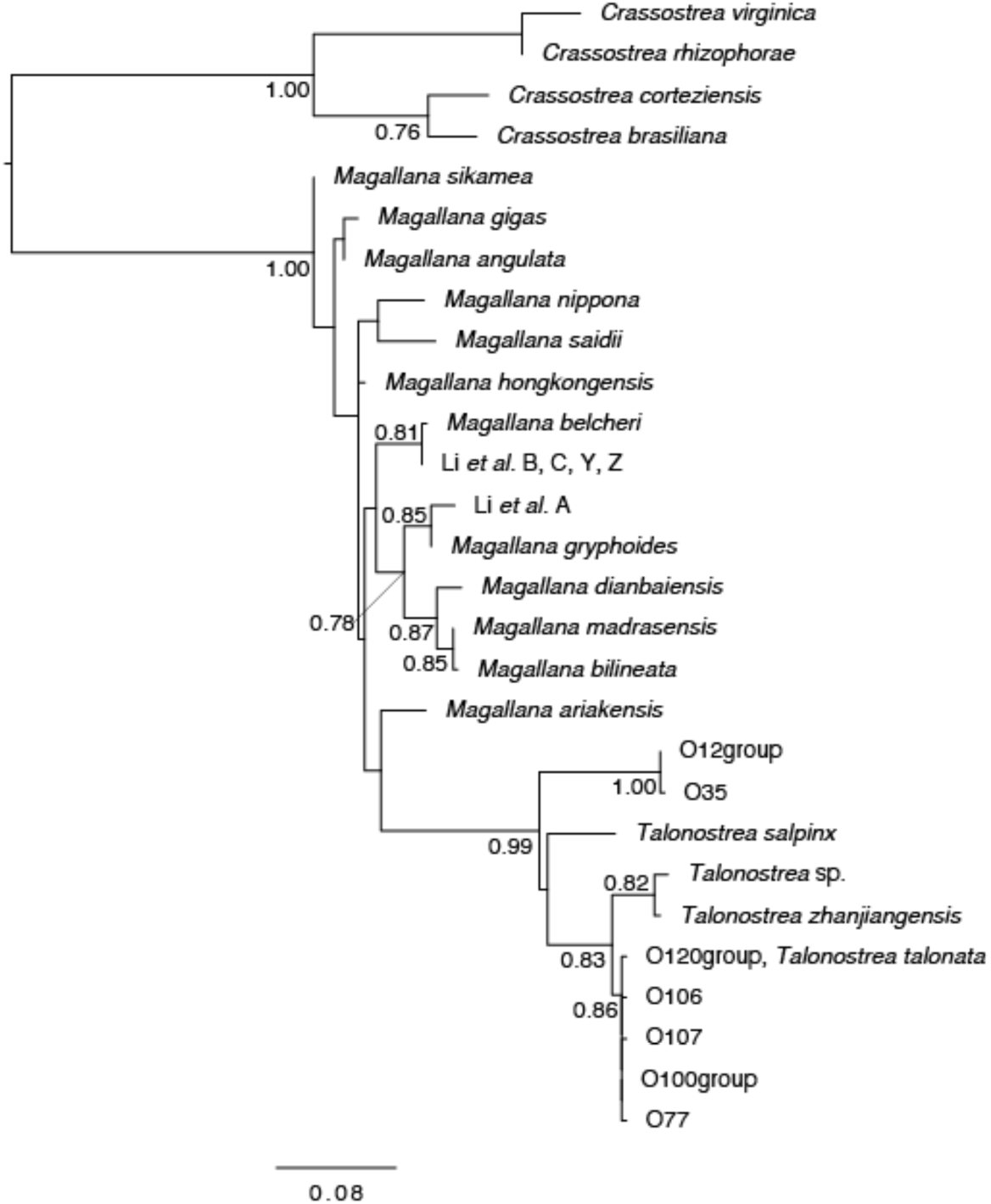
Maximum-likelihood tree for Crassostreinae oysters constructed using 16S rRNA gene sequence. Bootstrap values (≥0.70) are shown on nodes. Reference sequences of the genus *Crassostrea* were set as the outgroup. The scale bar is for the branch length and indicates the number of mutations per site.

**FIGURE 3.**
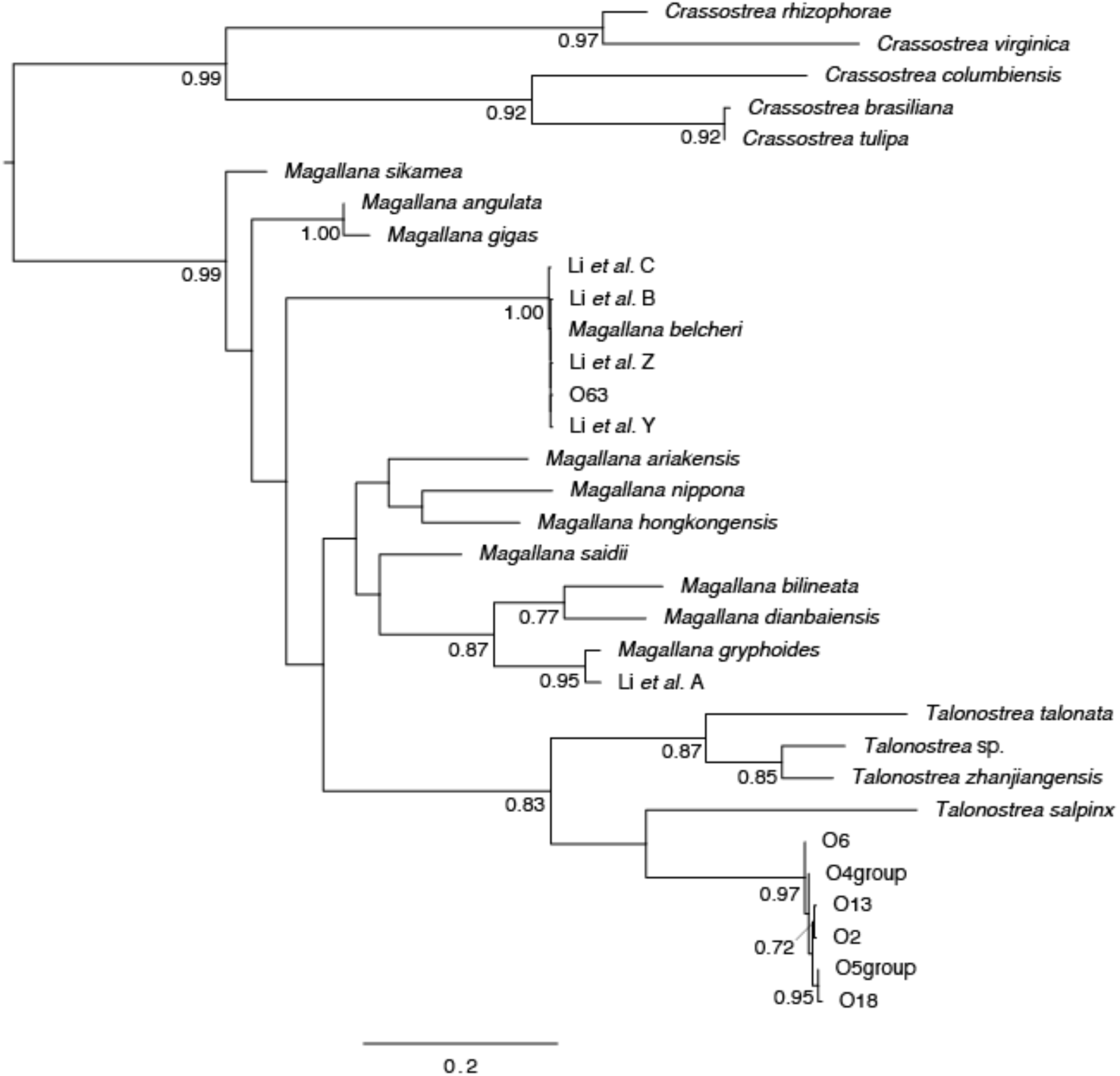
Maximum-likelihood tree for Crassostreinae oysters constructed using COI gene sequence. Bootstrap values (≥0.70) are shown on nodes. Reference sequences of the genus *Crassostrea* were set as the outgroup. The scale bar is for the branch length and indicates the number of mutations per site.

**Table 2.** Results of BLAST search.

| Sample origin | Taxa | COI | 16S |
| --- | --- | --- | --- |
| Mangrove root | Crassostreinae | 18 | 17 |
|  | *Saccostrea* | 18 | 24 |
| Cultured | Crassostreinae | 16 | 50 |
|  | *Saccostrea* | 7 | 22 |
| Spat | Crassostreinae | 2 | 13 |
|  | *Saccostrea* | 0 | 0 |
| Total | Crassostreinae | 36 | 80 |
|  | *Saccostrea* | 25 | 46 |

**Table 3.**
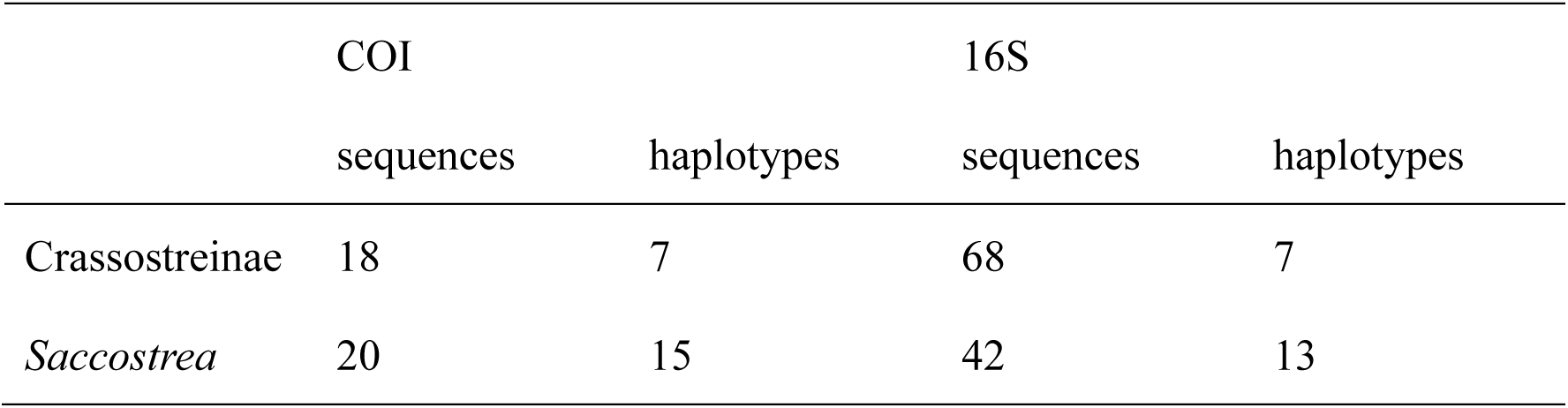
Number of sequences and haplotypes used for phylogenetic analyses.

|  | COI |  | 16S |  |
| --- | --- | --- | --- | --- |
|  | sequences | haplotypes | sequences | haplotypes |
| Crassostreinae | 18 | 7 | 68 | 7 |
| *Saccostrea* | 20 | 15 | 42 | 13 |

Among the 13 haplotypes of the 16S sequences of *Saccostrea* oysters, eight haplotypes representing 17 specimens were in the same clade as *S.* non-*mordax* lineage B (Figure 4). A single haplotype representing two specimens formed a clade with *S.* non-*mordax* lineage D. High-quality COI sequences were not retrieved from these two specimens. The remaining four haplotypes, representing 23 specimens, formed a clade with *S.* non-*mordax* lineage F. One haplotype derived from the four specimens was identical to the *S.* non-*mordax* lineage F reference sequence. Among the 15 haplotypes of the COI sequences of *Saccostrea* oysters, 3 haplotypes representing four specimens formed a clade with the *S.* non-*mordax* lineage F reference sequence (Figure 5). The remaining 12 haplotypes representing 16 specimens were classified into the same clade as the *S.* non-*mordax* lineage B reference sequence. There was one specimen in which the 16S sequence was identical to an *S*. non-*mordax* lineage F reference sequence. However, its COI sequence was identical to that of another specimen, which was classified as *S.* non-*mordax* lineage B. The specimen was unidentified.

**FIGURE 4.**
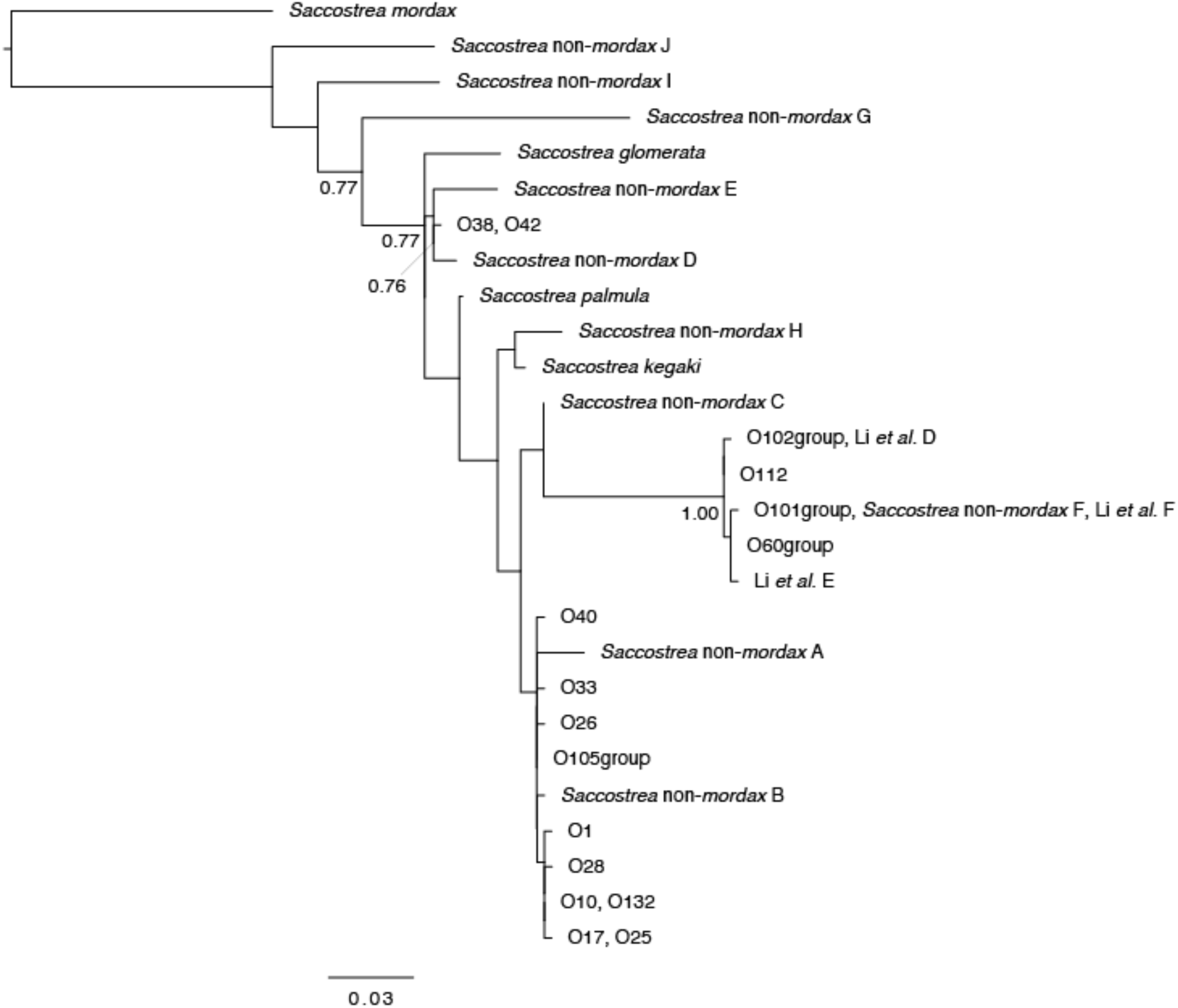
Maximum-likelihood tree for *Saccostrea* oysters constructed using 16S rRNA gene sequence. Bootstrap values (≥0.70) are shown on nodes. A reference sequences of *S. mordax* were set as the outgroup. The scale bar is for the branch length and indicates the number of mutations per site.

**FIGURE 5.**
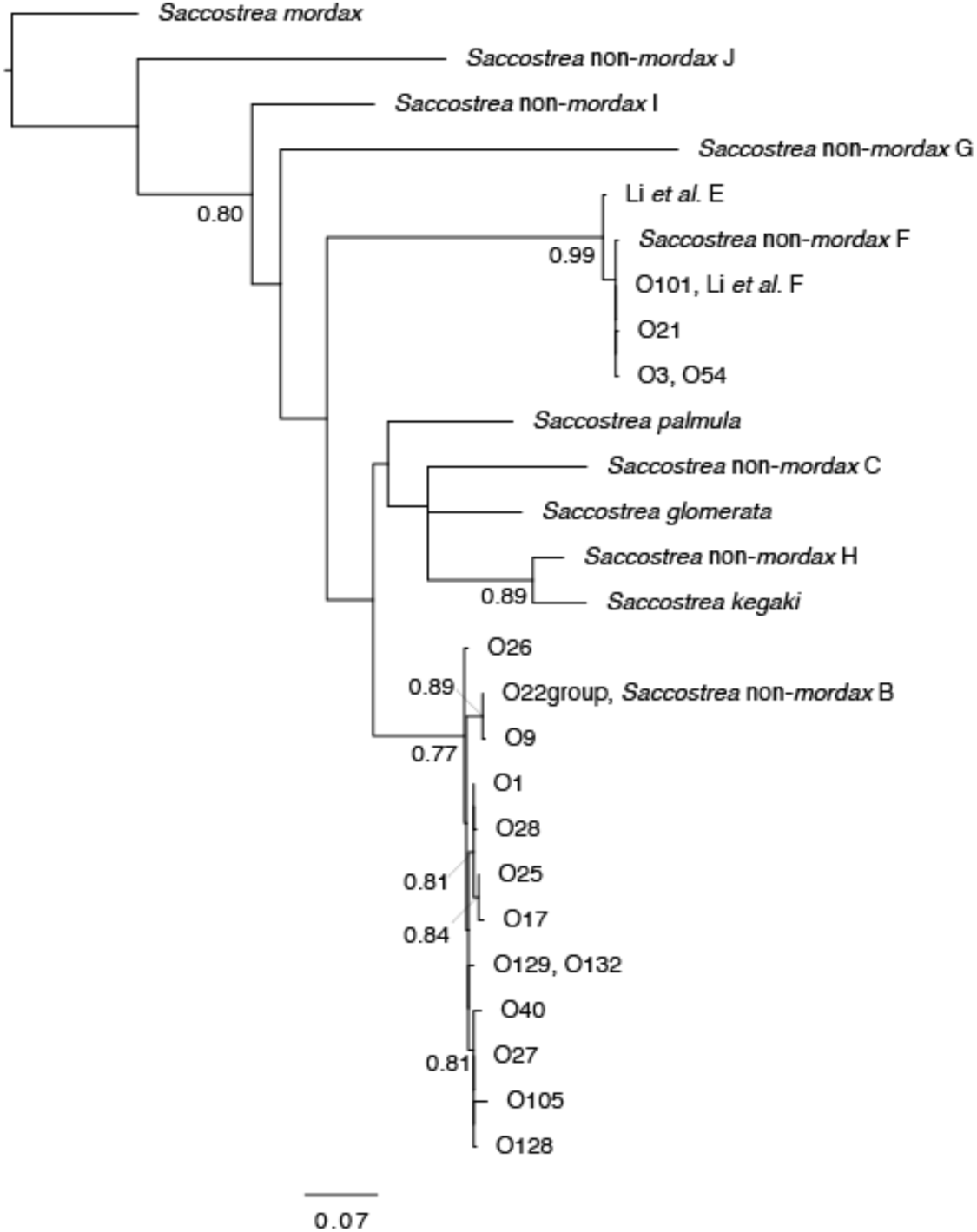
Maximum-likelihood tree for *Saccostrea* oysters constructed using COI gene sequence. Bootstrap values (≥0.70) are shown on nodes. A reference sequence of *S. mordax* were set as the outgroup. The scale bar is for the branch length and indicates the number of mutations per site.

Specimens identified from 16S or COI were counted separately based on the origin of the sample (Table 4). Four operational taxonomic units (OTUs) were identified in both the mangrove root and cultured oysters in Pa-yi-kyun village, but their compositions differed. *Talonostrea* sp.1 was discovered only in the mangrove root, whereas *T. talonata* was only derived from cultured oysters. Only sequences of *T. talonata* were retrieved and identified from the settled spat.

**Table 4.** Identity of oysters inferred from ML tree of COI or 16S.

| Sample origin | Taxa | COI | 16S | COI or 16S |
| --- | --- | --- | --- | --- |
| Mangrove root | *Talonostrea* sp.1 | 17 | 17 | 18 |
|  | *Saccostrea non-mordax* B | 11 | 14 | 14 |
|  | *S. non-mordax* D | 0 | 2 | 2 |
|  | *S. non-mordax* F | 2 | 6 | 6 |
| Cultured | *Magallana belcheri* | 1 | 0 | 1 |
|  | *T. talonata* | 0 | 39 | 39 |
|  | *S. non-mordax* B | 4 | 3 | 4 |
|  | *S. non-mordax* F | 2 | 16 | 16 |
| Spat | *T. talonata* | 0 | 12 | 12 |

**Table 5.** Shell morphology. Presence or absence of chomata, dark stain on the interior valve, radial ribs of the left valve, and a black or white steak on the outer surface of the right valve. ND indicates not determined because of fouling or cement on shell surface.

| Taxa | No. | Chomata |  | Stain |  | Radial rib |  |  | Streak |  |  |
| --- | --- | --- | --- | --- | --- | --- | --- | --- | --- | --- | --- |
|  |  | + | - | + | - | + | - | ND | + | - | ND |
| *Talonostrea* sp.1 | 2 | 0 | 2 | 0 | 2 | 0 | 2 | 0 | 0 | 2 | 0 |
| *T. talonata* | 25 | 0 | 25 | 2 | 23 | 22 | 0 | 3 | 9 | 12 | 4 |
| *Magallana belcheri* | 1 | 0 | 1 | 0 | 1 | 0 | 0 | 1 | 0 | 0 | 1 |
| *Saccostrea non-mordax* B | 4 | 4 | 0 | 2 | 2 | 0 | 4 | 0 | 0 | 0 | 4 |
| *S. non-mordax* F | 13 | 13 | 0 | 11 | 2 | 2 | 3 | 8 | 7 | 5 | 1 |

### Shell morphology

The shell characteristics of each taxon are summarised in Table 3. Three-D stacked images of the shells of nine specimens are shown in Figure 6. Most of the valves of the mangrove root specimens were not stored in good condition after DNA analysis. As a result, the shell morphology of *S.* non-*mordax* lineage D was not observed. *Talonostrea* sp. 1 (Figure 6A1–4, B1–4): There were no chomata. There was no dark staining on the inner surface of the valves; however, they had a slightly yellowish, dirty colour. The left valve did not have radial ribs. One of the two specimens exhibited four-pointed protrusions on the left valve. In the other specimen the left valve did not protrude. The marginal part of the left valve was extended like a flap, and the left valve was considerably longer than the smaller, flat right valve. No streaks were visible on the outer surface of the right valve. *T. talonata* (Figure 6C1–4, D1–4): The left valves had radial ribs. There were no chomata. In most cases (23 out of 25), there was no dark staining on the internal surface of the valves. The left valves had radial ribs in all specimens except in cases where the surface of the left valve was not visible because of the attached substrates. A considerable number of specimens (9 out of 21 observable specimens) had white or dark streaks on the surface of the right valve.

**FIGURE 6.**
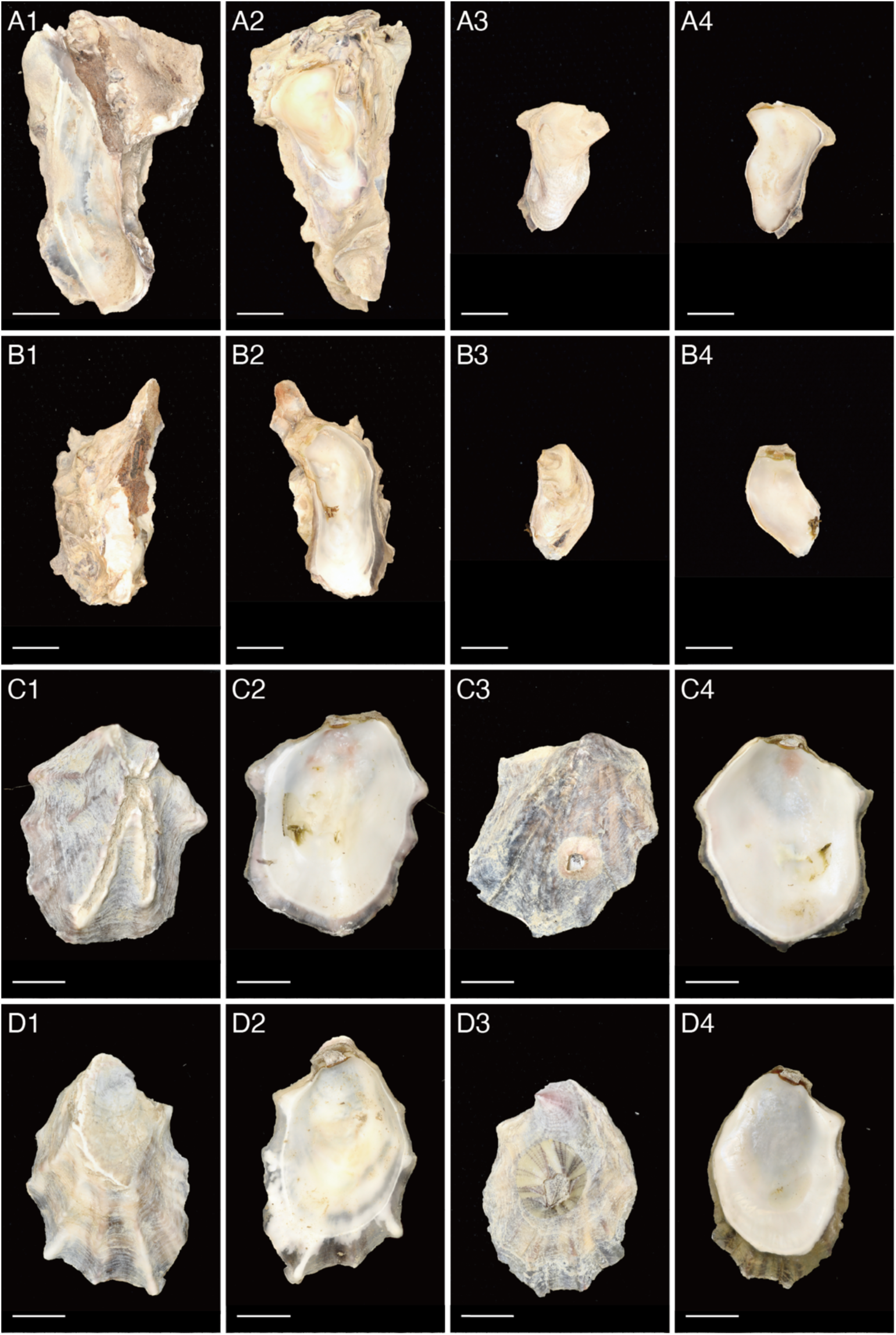
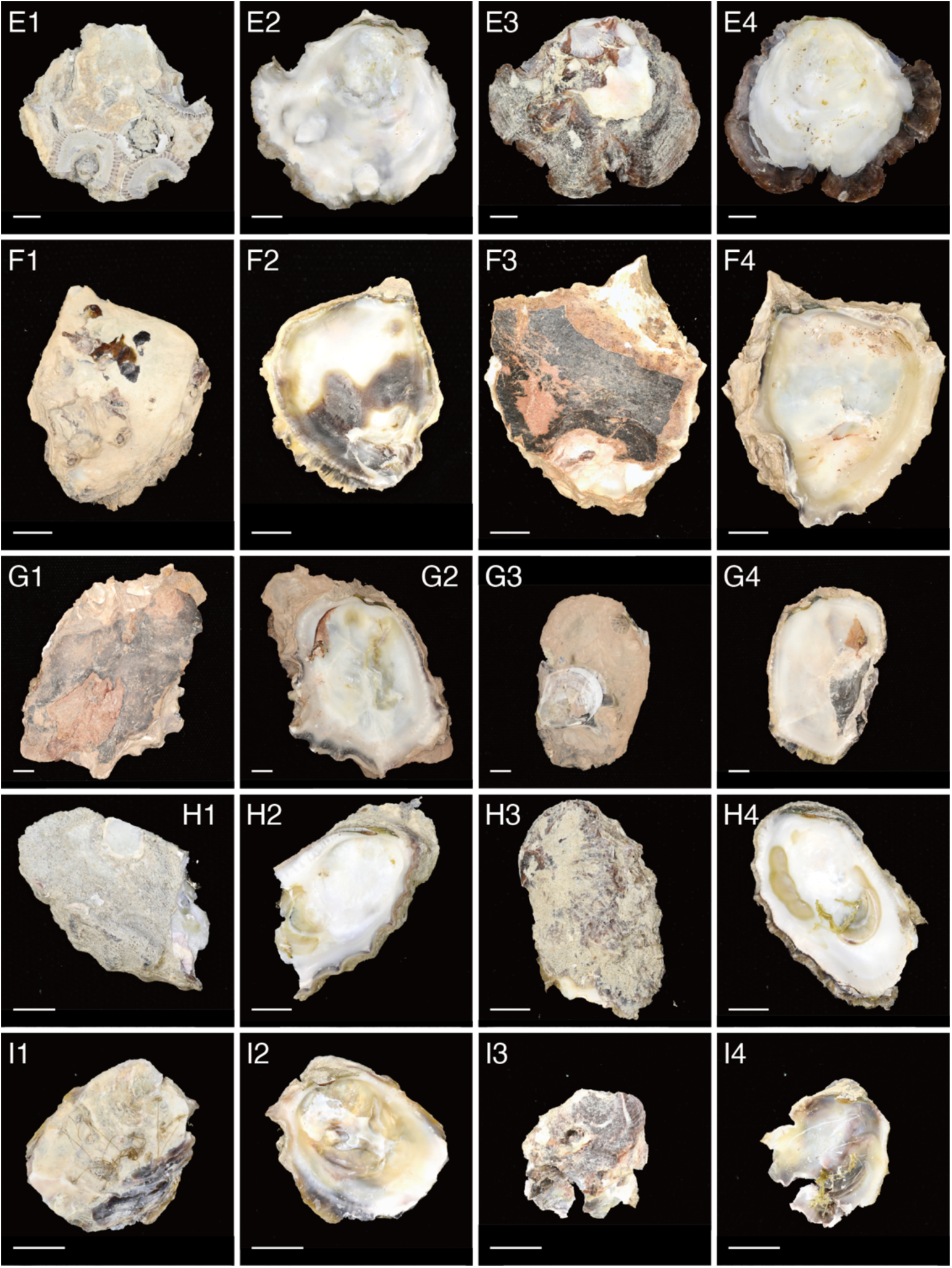
Shells of Crassostreinae and *Saccostrea* oysters. **A1 – 4**, O18, *Talonostrea* sp. 1, **B1 – 4**, O19, *Talonostrea* sp. 1, **C1 – 4**, O93, *Talonostrea talonata*, **D1 – 4**, O94, *T. talonata,* **E1 –4**, O63, *Magallana belcheri*, **F1 – 4**, O17, *Saccostrea* non-*mordax* lineage B, **G1 – 4**, O30, *Saccostrea* non-*mordax* lineage B, **H1 – 4**, O101, *Saccostrea* non-*mordax* lineage F, **I1 – 4**, O109, Saccostrea non-mordax lineage F, **A – I1**, left shell outside. **A – I2**, left shell inside. **A – I3**, right shell outside. **A – I4**, right shell inside. Scale bar = 5 mm.

*Magallana belcheri* (Figure 6E1–4): Only one specimen was examined in the present study. No chomata or staining was observed on the internal surface of the valves. The surfaces of the left and right valves were covered with substance; therefore, no radial ribs or streaks were observed.

*Saccostrea* oysters (Figure 6F1–4, G1–4: non-*mordax* lineage B; H1–4, I1–4: non- *mordax* lineage F): All *Saccostrea* oysters had chomata on the internal surfaces of the valves. Dark staining of the internal valve was observed in a high percentage (50% in non-*mordax* lineage B and 85% in non-*mordax* lineage F). No radial rib was present on the left valve of non-*mordax* lineage B; however, a radial rib was present in at least two specimens of non-*mordax* lineage F. Approximately half (7 out of 12) of the non- *mordax* lineage F specimens, in which the outer surface of the right valve was visible, had a white streak from the umbo to the shell margin.

## Discussion

In this study, *Talonostrea* oysters were recorded for the first time along the coast of Bengal Bay. *Talonostrea talonata* has been recorded from China, Japan, Peru, and Brazil (Li and Qi 1994; Inaba and Torigoe 2004; Li et al. 2017b; Cavaleiro et al. 2019). Also, *Talonostrea salpinx* was first described in the Persian Gulf in 2021 (Al-Kandari et al. 2021). Recently, another undescribed *Talonostrea* sp. was recorded in Australia (McDougall et al. 2024; Wells et al. 2024). Therefore, although records are scarce, the genus *Talonostrea* may be distributed throughout the coast of the Indo-Pacific and South Atlantic Oceans. However, we did not examine the anatomical details of these specimens. It is necessary to examine the anatomical details of the gills and digestive tract to compare the taxonomic characteristics of the previously described species (Li and Qi 1994; Al-Kandari et al. 2021). Therefore, we refrained from describing a new species and tentatively retained *Talonostrea* sp. 1, which is genetically separated from other species of the genus *Talonostrea*.

Three lineages of *Saccostrea* non-*mordax* (B, D, and F) were identified. It is difficult to distinguish these lineages based on shell morphology, as stated by Sekino and Yamashita (2016). A previous taxonomic study on oysters from the Taninthary Region of Myanmar reported a single lineage, F (*S. malabonensis*) (Li et al. 2017a). Our study added two lineages (B and D) from the very limited sampling; thus, three out of ten lineages that Sekino and Yamashita (2016) defined are now recorded from Myanmar. As the number of samples increases, more lineages will be recorded.

According to past research findings in Myanmar, Thi-Thi-Lay (1983, p.11) noted in her description of *S. cucullata* (Born) that a conspicuous streak of dark purple, black, or white colour is generally present on the right valve and runs from the hinge line towards the outer margin. She noted that most of the spat collected in both areas had one white or black stripe from the hinge to the shell edges. This proved that most of the spat settling was *S. cucullata* (Thi-Thi-Lay 1983, p.40). In the present study, a considerable proportion of *T. talonata* had white or dark streaks on the surface of the right valve. This suggested that *T. talonata* could have been present in the samples observed by Thi-Thi-Lay. Owing to the increasing progress of studies in Myanmar, the distribution of *Talonostrea* spp. in Myanmar is expected to be revealed in the near future.

All the specimens examined in this study belonged to Crassostreinae or *Saccostrea*. Most Crassostreinae were not *M. belcheri* but *Talonostrea* spp. Among the 126 successfully analysed specimens, a single specimen was identified as *M. belcheri*. This suggested the difficulty in establishing a culture of the larger edible tropical oyster *M. belcheri*, which depends only on natural spat collection from the wild. To obtain spat from the wild, further knowledge about the settlement of *M. belcheri* in this region, such as the peak settling season, preferred depths, and substrate, is necessary. However, until a simple method to distinguish spat of *M. belcheri* from spat of other species is established, artificial seedlings of the spat would be a more realistic option, because the method has already been established (Tanyaros 2011; Tanyaros and Tarangkoon 2016). In Myeik, research on oyster seed production from hatcheries has been conducted since last year, and positive results have been achieved. Continued research promises the establishment of an oyster culture industry in Myanmar shortly.

## Acknowledgements

We thank Prof. Nyo-Nyo-Tun of Mawlamyine University for her efforts in arranging a cooperative study on oyster aquaculture between Myeik University and JIRCAS, field sampling and obtaining permission to bring genetic samples back to Japan, while serving as the head of the Marine Science Department at Myeik University. We thank the staff of the Marine Science Department, Myeik University, and the local assistants, especially Dr. Aung-Aung-Aye, Ms. Ei-Thal-Phyu, and Mr. Myokoko-Thwin, for their assistance with sampling. We sincerely thank the Department of Fisheries, Ministry of Agriculture, Livestock and Irrigation, the Republic of the Union of Myanmar, especially Dr. Aung-Naing-Oo, Director of the Aquaculture Division, for coordinating permission procedures for sampling in Myanmar and material transfer to Japan. We also thank Dr. Saito Hajime of JIRCAS (present affiliation: FRA) for his assistance in the preparation of oyster samples and Dr. Marcy N. Wilder of JIRCAS for her support with DNA analysis. We would like to thank Editage (www.editage.jp) for English language editing. This research was mainly conducted using a grant budget from the Japan International Research Centre for Agricultural Sciences and partially supported by Japanese Grants-in-Aid for Scientific Research (No: 20KK0141).

## Author contributions

M.T. and T.Y. contributed equally to this article as the main researchers. T.Y. and B.J.K. organised and measured the oyster samples, conducted DNA extraction and PCR operations, and analysed the nucleotide sequences. M.T. and Z.S. performed phylogenetic analysis of the data and took morphological photographs of the shells. C.A. provided local fishery information and managed the study. M.T. prepared the draft manuscript, and all authors revised and approved the final manuscript.

